# Impact of cell culture model systems on SARS-CoV-2 and MERS-CoV infection dynamics and antiviral responses

**DOI:** 10.1101/2025.02.24.639954

**Authors:** Kim Chiok, Mark Fenton, Darryl Falzarano, Neeraj Dhar, Arinjay Banerjee

## Abstract

Cell cultures are widely used to study infectious respiratory diseases and to test therapeutics; however, they do not faithfully recapitulate the architecture and complexity of the human respiratory tract. Lung organoids have emerged as an alternative model that partially overcomes this key disadvantage. Lung organoids can be cultured in various formats that offer potential for studying highly pathogenic viruses. However, the effects of these different formats on virus infection remain unexplored, leaving their relative value unclear. In this study, we generated primary lung organoids from human donor cells and used them to derive monolayers and air-liquid interface (ALI) cultures with the goal of comparing the replication kinetics of two circulating highly pathogenic coronaviruses, severe acute respiratory syndrome coronavirus 2 (SARS-CoV-2) and Middle East respiratory syndrome coronavirus (MERS-CoV). Infection studies revealed that organoid-derived monolayers displayed limited infection and the innate immune response was impaired against bacterial lipopolysaccharide (LPS) but not against virus-like double-stranded dsRNA or poly(I:C). Meanwhile, organoids and organoid-derived ALI cultures retained viral permissivity, with ALI cultures displaying diverse antiviral immune responses against both coronaviruses. SARS-CoV-2 and MERS-CoV demonstrated differential replication kinetics in organoid and organoid-derived ALI cultures. Therefore, primary organoid-derived cells in two-dimensional monolayer or three-dimensional ALI formats influence virus infection and host antiviral responses. Our study informs the selection of culture conditions for organoid-based respiratory disease research and therapeutic testing.

**Importance:** The COVID-19 pandemic heralded the upsurge in human-derived lung organoid based studies due to their cellular heterogeneity that emulates the cellular complexity of the respiratory tract. A major disadvantage of organoid models resides in their apical-in conformation that “hides” cells and proteins that are typically exposed to the air-liquid interface (ALI) in the airways and are targets of viruses. Here, we generated monolayers and ALI cultures to facilitate cell exposure to highly relevant pathogens and compared them to parental organoids. Organoids at the ALI captured infection and immune responses better than organoids and organoid-derived monolayer cultures. Organoids at the ALI are a viable approach to improve identification and characterization of virus infection, host responses, and therapeutic testing.

## 1. Introduction

Traditionally, researchers have relied on two-dimensional (2D) cell culture models such as immortalized cell lines to uncover molecular mechanisms of disease, understand the host immune response against infectious viral agents, identify therapeutic targets, and assess drug efficacy (1). Even for studying respiratory viruses, cells are typically cultured as 2D monolayers, submerged in nutrient rich media that supports their growth and replication. While these cell lines are important tools for studying disease mechanisms, they often fail to replicate the cellular heterogeneity, three-dimensional architecture, and physiological responses of respiratory tissue, which is naturally exposed to environmental air in the airways. Animal models are useful to validate *in vitro* observations, however, these models can be ethically challenging and yield inconclusive outcomes due to anatomical and cellular differences when compared to humans. This is a major bottleneck when studying newly emerging viruses where time is critical and the absence of validated animal models impedes the evaluation of therapeutics (2). As emerging pathogens are a continuous threat to global health, human-like *in vitro* models are key to identify, develop, and test prophylactic and therapeutic measures.

Organoids derived from human cells have emerged as an alternate model that recapitulates the structural and compositional complexity of the native organ. Organoids are composed of self-renewing stem cells that differentiate into various cell types present in the tissue of origin and self-assemble into three-dimensional (3D) microtissues (3). Adult <u>s</u>tem <u>c</u>ells (ASC) from donor tissue are viable sources for organoid development without the lengthy differentiation process used for induced pluripotent stem cells (iPSCs) or the oncogenic mutations of cancer derived organoids (4). Donor-derived organoids preserve the individual-level diversity that influences immune response, susceptibility to pathogens, metabolism and tolerance to pharmaceuticals. These advantages have poised organoids as promising approaches for use in preclinical therapeutic screening and has received authorization from the US Food and Drug Administration (FDA) for this purpose (5, 6). Human respiratory organoids are being actively used to study respiratory pathogens like Influenza A Virus (IAV) (7, 8), Respiratory Syncytial Virus (RSV) (9), and Human Adenovirus type 3 and type 55 (10), and the recently emerged coronaviruses severe acute respiratory syndrome coronavirus 2 (SARS-CoV-2) and Middle East respiratory syndrome coronavirus (MERS-CoV) (11, 12). Despite this progress, studies comparing 2D and 3D cell culture models and their impact on SARS-CoV-2 and MERS-CoV replication are limited.

SARS-CoV-2 emerged in late 2019 to cause the COVID-19 (coronavirus disease 2019) pandemic. As of December 2024, the World Health Organization (WHO) has reported over 7 million deaths worldwide since the COVID-19 pandemic started, which is likely an underestimate of the true impact of this pandemic (13). COVID-19 mortality rates are estimated to be between 0.1 to 5% (14). The WHO data show that the virus continues to disseminate (15), propelled by the emergence of viral variants that can at least partially evade protection from existing vaccines (16). MERS-CoV emerged in 2012 and continues to cause outbreaks of severe viral pneumonia with about 35% case fatality rate (17). MERS-CoV remains a pathogen of concern and a pandemic threat. Indeed, research into SARS-CoV-2 and MERS-CoV can provide insights into coronavirus biology which will inform the development of prophylactic and therapeutic interventions. Human-derived experimental models are thus essential to understand and mitigate risks posed by emerging coronaviruses and other respiratory pathogens.

In this study, we established human donor-derived lung organoids from which we derived traditional cell monolayers and air-liquid interface (ALI) cultures and compared infectivity using currently circulating highly pathogenic coronaviruses, SARS-CoV-2 and MERS-CoV. Lung organoids and derived monolayers and ALI cultures showed distinct differences in virus infection and transcriptional regulation of antiviral genes upon immune stimulation and virus infection. While monolayers transitioned into a virus-resistant phenotype, ALI cultures sustained viral infection and antiviral response against virus infection. Despite their common origin, differences between two-(2D) and three-dimensional (3D) cultures emphasize the need for careful selection of cell culture models for respiratory infectious disease studies and therapeutic testing.

## 2. Results

### 2.1 Cell culture platforms influence organoid growth properties

We obtained lung tissue sample from a donor patient and used previously established protocols (21) to generate human lung organoids (hLO). These hLO were then cultured and maintained under three different formats – traditional two-dimensional adherent monolayers (hLOm); as three-dimensional air-liquid interface cultures (hLO ALI) or passaged as three-dimensional lung organoids (hLO) (**Fig. 1A**).

**Figure 1.**
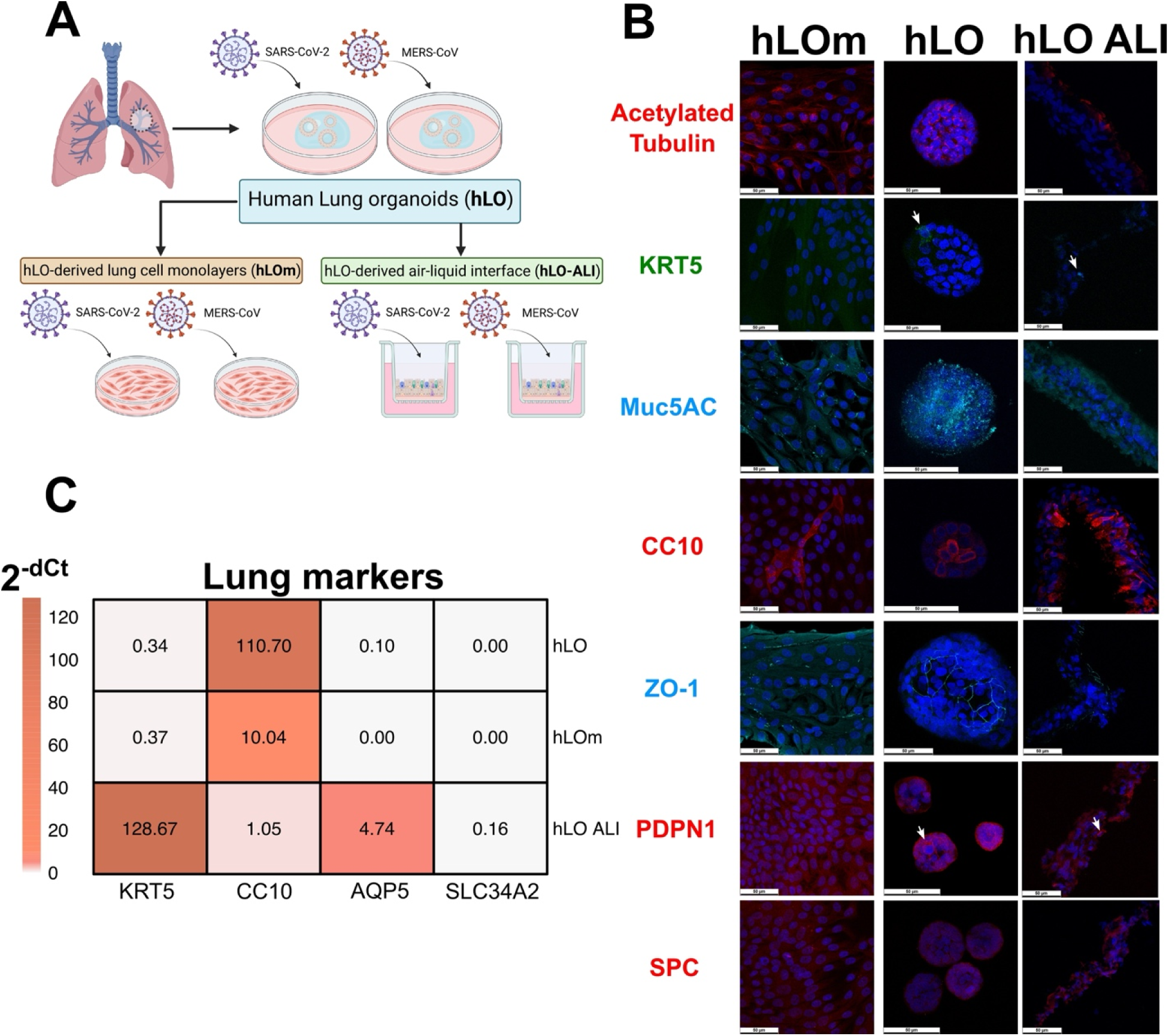
Generation of donor-derived lung organoids (hLO), organoid-derived monolayers (hLOm) and ALI (hLO ALI) cultures. (A) Schematic of hLO generation and derivation of hLOm and hLO-ALI cultures, followed by infection with SARS-CoV-2 or MERS-CoV. Schematic created using BioRender. (B) Identification of lung cell markers. hLOm, hLO and hLO ALI cells were fixed in 4% paraformaldehyde, permeabilized and stained with antibodies against acetylated tubulin (cilia), KRT5 (Basal Stem Cells, white arrows), Muc5AC (mucus and mucus producing cells, Goblet Cells), CC10 (Club Cells), ZO-1 (zona occludens-1, tight junctions), PDPN1 (podoplanin-1, AT1 cells, white arrows), SPC (surfactant protein C, AT2 cells), and DAPI (nuclei, blue). Cells were imaged on a confocal microscope using a 63x objective. Scale bars correspond to 50 µm. (C) Gene expression of lung markers KRT5, CC10, AQP5 (aquaporin 5, AT-1 cells), and SLC34A2 (AT-2 cells) in hLO, hLOm and hLO ALI was determined by RT-qPCR assays. Samples were assayed in technical duplicates, dCt was normalized to GAPDH and expression was calculated by 2^-dCt^. The mean of three independent samples is presented, error bars are standard error of the mean (SEM).

To characterize the distribution of different cell types in the hLOm and hLOs, we carried out immunofluorescence staining using antibodies directed to cell type specific markers **(Fig. 1B)**. Acetylated tubulin localized along cytoplasmic microtubules in hLOm, within the hLOs indicating an apical-in orientation, and consistent with cilia formation in the apical-out orientation in hLO ALI cultures (**Fig. 1B**). We identified KRT5+ cells (basal stem cells), the mucus component Muc5AC produced by Goblet cells (22), the tight junction protein zona occludens-1 (ZO-1), and the alveolar type 1 (AT1) protein podoplanin-1 (PDPN1) in hLO and hLO ALI cultures identified as individual cells stained with corresponding markers. Club cells (CC10) typical of bronchiolar epithelium (23) were more abundant in hLOs and hLO ALI relative to hLO monolayers. In hLO ALI, CC10 cells appeared containing secretory granules that suggested functional production and secretion of surfactant (**Fig. 1B**). The absence of several markers in hLOm suggested that this format does not foster various cell types and structures typical of the lungs (**Fig. 1B**). We did not identify specific individual cells positive for Surfactant Protein C (SPC) in any of the cultures, suggesting limited presence of alveolar type 2 (AT2) cells (**Fig. 1B**).

We also employed RT-qPCR assays to further confirm the composition of hLOs and the derived culture models (**Fig. 1C**). We found that transcript levels of KRT5 (marker for basal stem cells) were higher in hLO ALI, consistent with ALI promotion of basal stem cell growth reported previously (24). CC10 transcript levels were higher in hLO relative to hLOm and ALI cultures (hLO ALI). We determined transcription levels of AQP5 and SLC34A2 as additional markers of AT1 and AT2 cells, respectively by RT-qPCR. hLO ALI and hLO both had detectable levels of AT1-specific AQP-5 transcripts, particularly the ALI format which also had detectable levels of AT2-specific SLC34A2 (**Fig. 1C**). Therefore, our data demonstrate that cellular composition was different between hLO, hLOm and hLO ALI cultures derived from the same parental organoids. Our results suggest that 3D formats like hLO and hLO ALI consist of a mixture of lung airway and alveolar cells capable of producing surfactant and form permeability barrier (25–27) that are absent in cell culture monolayers.

### 2.2. Culture format influences innate immune response against virus- and bacteria-like stimuli

Next, we examined the cellular response against bacteria-like and virus-like stimuli in hLOm and hLO as prototypic 2D and 3D culture systems. hLOm and hLO were transfected with multiple doses of the dsRNA analog polyinosinic-polycytidylic acid [poly(I:C)] to mimic viral infection. Poly(I:C) was delivered successfully in both hLOm and hLO as observed by live cell imaging (**Fig. 2A**). hLOm and hLO responded to this stimulus by upregulating the transcripts for antiviral genes like interferon-beta (*IFNβ*) (**Fig. 2B**), interferon Induced Protein with tetratricopeptide 1 (*IFIT1*) (**Fig. 2C**), 2’-5’-oligoadenylate synthetase 1 (*OAS1*) (**Fig. 2D**) and MX Dynamin Like GTPase 1 (*MX1*) (**Fig. 2E**). While most responses were comparable between hLO and hLOm, upregulation of *IFNβ* was nearly 100-fold higher in hLOm (**Fig. 2B**). We (28) and others (29) previously identified *IFNβ* and this set of interferon stimulated <u>g</u>enes (ISGs) as relevant for antiviral responses against coronaviruses. Therefore, both hLO and hLOm upregulate antiviral genes upon encountering intracellular virus-like stimuli.

**Figure 2.**
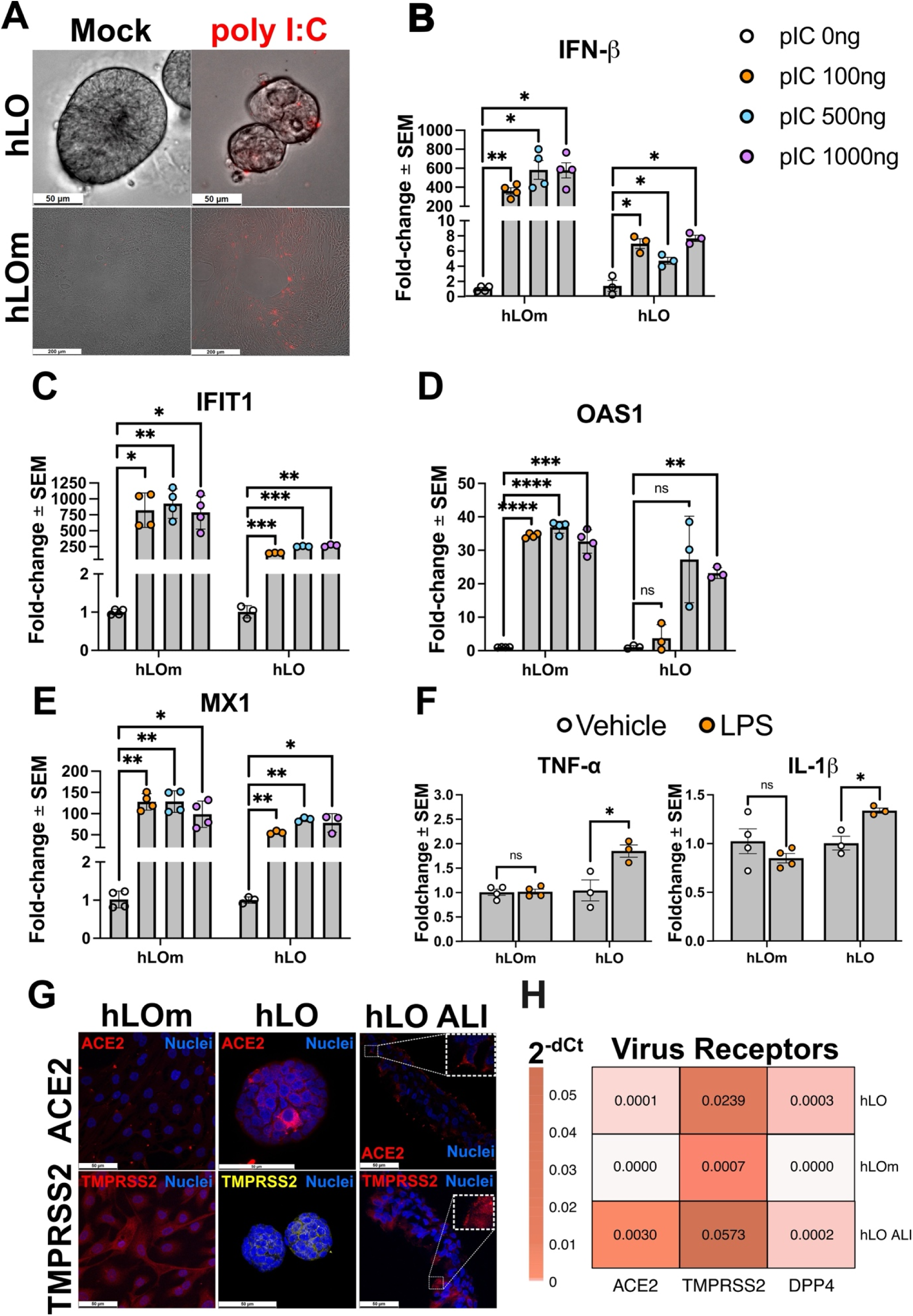
Evaluation of immunocompetence against virus- and bacteria-like stimuli. (A) hLO and hLOm were mock-transfected or transfected with 2µg of rhodamine labeled polyI:C (red). Live cells were imaged at 48 hours post transfection. Scale bars correspond to 50 µm (hLO, top panels) and 200 µm (hLOm, bottom panels). (B) hLO and hLOm were transfected with increasing doses of polyI:C (pIC) for 48 hours. Total RNA was collected for RT-qPCR assays to determine gene expression of *IFNβ* (B)*, IFIT1* (C), *OAS1* (D), and *MX1* (E). (F) hLO and hLOm were treated with either vehicle (LPS-free water) or LPS (100ng/mL) for 6 hours. RNA was collected for RT-qPCR determination of *TNF-α* and *IL-1β* transcript levels. (G) hLOm, hLO and hLO ALI were fixed in paraformaldehyde 4%, permeabilized and stained with antibodies against ACE2 and TMPRSS2, and counterstained with DAPI (nuclei, blue). Cells were imaged using a confocal microscope at immersion (63x). Individual positive cells are presented as insets highlighted with white boxes. Scale bars correspond to 50 µm. (I) Gene expression levels of *ACE2*, *TMPRSS2* and *DPP4* in hLO, hLOm and hLO ALI was determined by RT-qPCR assays. Samples were assayed in technical duplicates, dCt was normalized to *GAPDH* and expression was calculated by 2^-dCt^. The mean of three independent samples is presented. For RT-qPCR assays in B to F, samples were assayed in technical duplicates, dCt was normalized to GAPDH and expression was calculated as fold-change relative to vehicle treated controls by 2^-ddCt^ method. The mean of three independent samples is presented, error bars are standard error of the mean (SEM). Statistical analysis was performed by One-way ANOVA (B-E) or Student’s T Test (F). *, *P* < 0.05; **, *P* < 0.01; ***, *P* < 0.001; ****, *P* < 0.0001.

Lipopolysaccharide (LPS) is a major component of the outer membrane of gram-negative bacteria that induces production of tumor necrosis factor alpha (*TNF-α*) and interleukin 1 beta (*IL-1β*) to mediate inflammation and acute lung injury in cellular and animal models (30). hLOs exposed to LPS from pathogenic *E. coli* O111:B4 responded by upregulating the proinflammatory genes *TNF-α* and *IL-1β* (**Fig. 2F**). In contrast, hLOm did not upregulate these proinflammatory cytokines and remained unresponsive to LPS stimuli. These results indicate that donor-derived hLO and corresponding hLOm are immunocompetent and responsive to virus-like stimuli, but only hLOs respond to LPS. Thus, transition from 3D to 2D culture formats may influence the breadth of the innate immune response, with hLOm missing relevant pulmonary responses such as those directed against bacterial LPS.

Next, we aimed to map the distribution of coronavirus receptors in hLOm, hLO and hLO ALI. Immunofluorescence staining revealed that distinct cells within parental hLOs expressed angiotensin converting enzyme 2 (ACE2) and TMPRSS2, the cellular receptor for SARS-CoV-2 and its entry cellular co-factor, respectively (31) (**Fig. 2G**). We did not detect ACE2 in monolayers, whereas TMPRSS2 appeared less intense and diffuse in cytoplasms of cells in monolayers (**Fig. 2G**). Individual cells in hLO ALI also expressed ACE2 and TMPRSS2, the latter of which localized to the cell membrane and cilia (**Fig. 2G**), similar to what has been reported previously for primary cell nasal epithelium ALI cultures (32). Despite our best efforts, we were unable to detect the DPP4 protein, the receptor for MERS-CoV (33), by immunostaining. Additional RT-qPCR assays detected transcripts for ACE2 and TMPRSS2 predominantly in hLO and hLO ALI (**Fig. 2H**), with higher transcript levels in hLO ALI, which is consistent with our immunostaining findings. Meanwhile, we also detected *DPP4* transcripts in hLO and hLO ALI, but not in hLOm (**Fig. 2H**). These results suggest that in addition to differences in cellular composition and architecture, hLO cells under different culture formats may differ in their surface protein, such as viral receptor expression profiles and thus, susceptibility to coronavirus infection.

### 2.3. 2D and 3D formats dictate differences in SARS-CoV-2 infectivity

Since 2D and 3D cultures differed in cell composition, architecture and breadth of response against pathogen-like stimuli, we next investigated whether culture format also impacts infection with respiratory viruses. We first used SARS-CoV-2 due to its recent relevance for global health. Parental hLO, hLOm, and hLO ALI cultures were infected with SARS-CoV-2 and followed for up to 7 days. Brightfield images show that hLO monolayers appeared to produce surfactant observed as darker areas on top of monolayers composed of cells with epithelial-like morphology (**Fig. 3A**). Meanwhile, hLOs retained characteristic spherical morphology with presence of cyst-like cavities and absence of cilia on the apical aspect of the organoids (apical-in). hLO ALI displayed packed pseudostratified epithelium with presence of beating cilia (**Supplementary video 1**) that actively moved mucus atop the culture, giving the appearance of whorls when viewed from top down (**Fig. 3A**).

**Figure 3.**
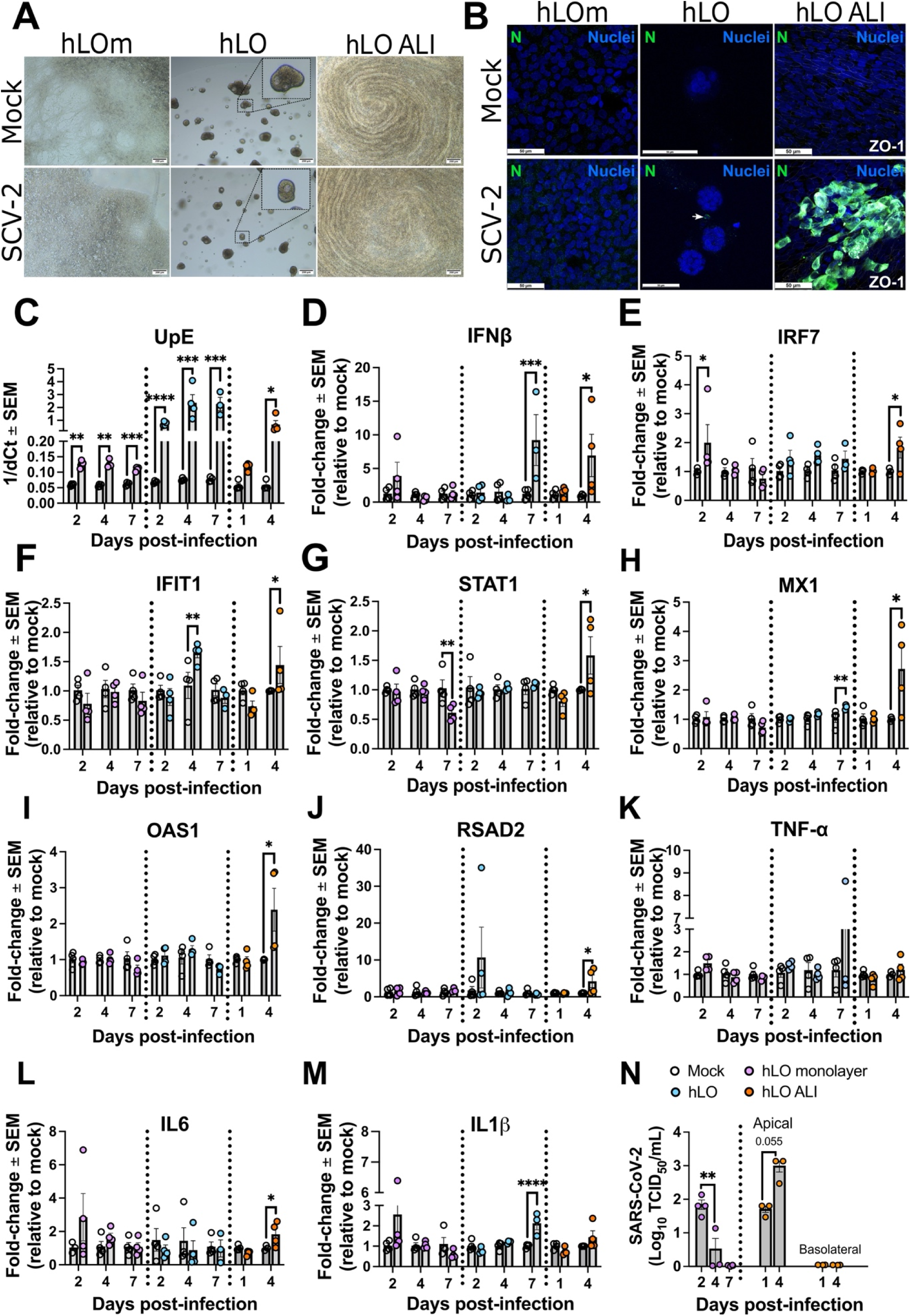
SARS-CoV-2 infectivity in donor-derived lung organoids (hLO), organoid-derived monolayers (hLOm) and ALI (hLO ALI) cultures. (A) Brightfield (5x) and (B) confocal imaging of immunostained cells mock infected or infected with SARS-CoV-2 (SCV2) at an MOI of 0.3 at 4 days post infection. Cells were fixed, permeabilized and stained with antibodies against the SARS-CoV-2 nucleoprotein (N) and DAPI (nuclei) and imaged at 63x. Scale bars correspond to 200 µm (A) and 50 µm (B). Total RNA was collected at specified times to determine expression of a viral genomic region upstream the E gene (UpE) (C) and host gene expression of *IFNβ* (D), *IRF7* (E), *IFIT1* (F), *STAT1* (G), *MX1* (H), *OAS1* (I), *RSAD2* (J), *TNF-α* (K), *IL6* (L) and *IL1*β (M) by RT-qPCR. (N) SARS-CoV-2 virus titers in supernatant of monolayers, and apical and basolateral aspects of ALI cultures by TCID_50_ assay. For RT-qPCR assays, samples were assayed in technical duplicates, dCt was normalized to GAPDH and virus gene expression was calculated as 1/dCt. Host gene expression was calculated as fold-change relative to mock controls by 2^-ddCt^ method. The mean of three independent samples is presented, error bars are standard error of the mean (SEM). Statistical analysis was performed by One-way ANOVA (C-M) or Student’s T Test (N). *, *P* < 0.05; **, *P* < 0.01; ***, *P* < 0.001; ****, *P* < 0.0001.

Despite the absence of apparent cytopathic effect (CPE) in all three types of culture upon SARS- CoV-2 infection (**Fig. 3A**), immunofluorescence staining confirmed the presence of the SARS- CoV-2 N protein in hLO and hLO ALI cultures (**Fig. 3B**). We evaluated tight junction status in ALI cultures by staining for the ZO-1 marker (**Fig. 3B**), which appeared unaffected by SARS- CoV-2 infection in our experimental conditions (**Fig. 3B**). We did not detect cells positive for SARS-CoV-2 N protein in hLOm (**Fig. 3B**). These observations were further confirmed by RT- qPCR assays which showed higher upregulation of viral genes (UpE) in hLO and hLO ALI cultures upon infection with SARS-CoV-2, when compared to hLOm (**Fig. 3C**) (28).

We then profiled the immune responses that were generated in the three culture systems upon SARS-CoV-2 infection. hLO and ALI cultures upregulated *IFNβ* transcripts at 7- and 4-days post-infection, respectively (**Fig. 3D**). SARS-CoV-2 infection led to the upregulation of several canonical antiviral ISGs in hLO ALI including IFN regulatory factor 7 (*IRF7*) (**Fig. 3E**), *IFIT1* (**Fig. 3F**), *STAT1* (**Fig. 3G**), *MX1* (**Fig. 3H**), *OAS1* (**Fig. 3I**), and Radical S-Adenosyl Methionine Domain Containing 2 (*RSAD2*) (**Fig. 3J**) at 4 days post infection. In case of hLO, SARS-CoV-2 infection led to upregulation of *IFIT1* (**Fig. 3F**), with the remaining ISGs failing to respond to virus infection. Analyses of key proinflammatory genes **(Fig. 3K-M)** demonstrated the upregulation of transcripts for Interleukin 6 (*IL6*) in hLO ALI at 4dpi (**Fig. 3L**), whereas transcripts for *IL1β* were upregulated in hLO at 7dpi (**Fig. 3M**). hLOm remained mostly unresponsive to virus infection and did not upregulate transcripts for our selected antiviral genes other than *IRF7* at 2dpi (**Fig. 3C**). Thus, hLO ALI cultures were more responsive to SARS-CoV- 2 infection, relative to parental hLO and hLOm.

hLOm and hLO ALI demonstrated the presence of SARS-CoV-2 via detection of viral transcripts (UpE, **Fig. 3C**). However, only ALI cultures had cells that stained positive for viral nucleoprotein (N) (**Fig. 3B**) and upregulated transcripts for antiviral and proinflammatory genes in response to SARS-CoV-2 **(Fig. 3D-M)**. We performed TCID_50_ assays to determine whether hLOm or hLO ALI can produce infectious virus progeny given that virus UpE levels were lower in these formats relative to hLO. Infected hLO ALI produced increasing amounts of infectious virus at the apical side of the ALI. In hLOm, virus was detected on Day 2 post-infection which then decreased and became undetectable by 4 days post infection (**Fig. 3N**). We did not detect infectious virus in ALI basolateral medium (**Fig. 3N**). These results show that despite their common origin, culture format impacts SARS-CoV-2 infection kinetics and immune response in primary donor lung organoid cells.

### 2.4. 2D and 3D formats dictate differences in MERS-CoV infectivity

We next investigated whether culture format also impacts MERS-CoV infection kinetics and host response. Like SARS-CoV-2, we did not observe CPE under bright field microscopy in either hLOm, hLO, or hLO ALI infected with MERS-CoV (**Fig. 4A**). Both hLO and hLO ALI had cells that stained positive for MERS-CoV nucleoprotein (N), whereas only few cells were positive for MERS-CoV N in hLOm (**Fig. 4B**). ALI cultures infected with MERS-CoV had areas of cell detachment, leaving gaps lined by infected cells and exposed cells in the basement layer that appeared as plaque-like lesions (**Fig. 4B**). These disrupted areas had lost the ZO-1 marker, suggesting loss of tight junctions and permeability barrier (**Fig. 4B**). RT-qPCR assays confirmed upregulation of viral gene transcripts (UpE) in hLOm, hLO and hLO ALI cultures (**Fig. 4C**). However, hLOm and hLO ALI cultures did not show time-dependent increase of UpE (**Fig. 4C**).

**Figure 4.**
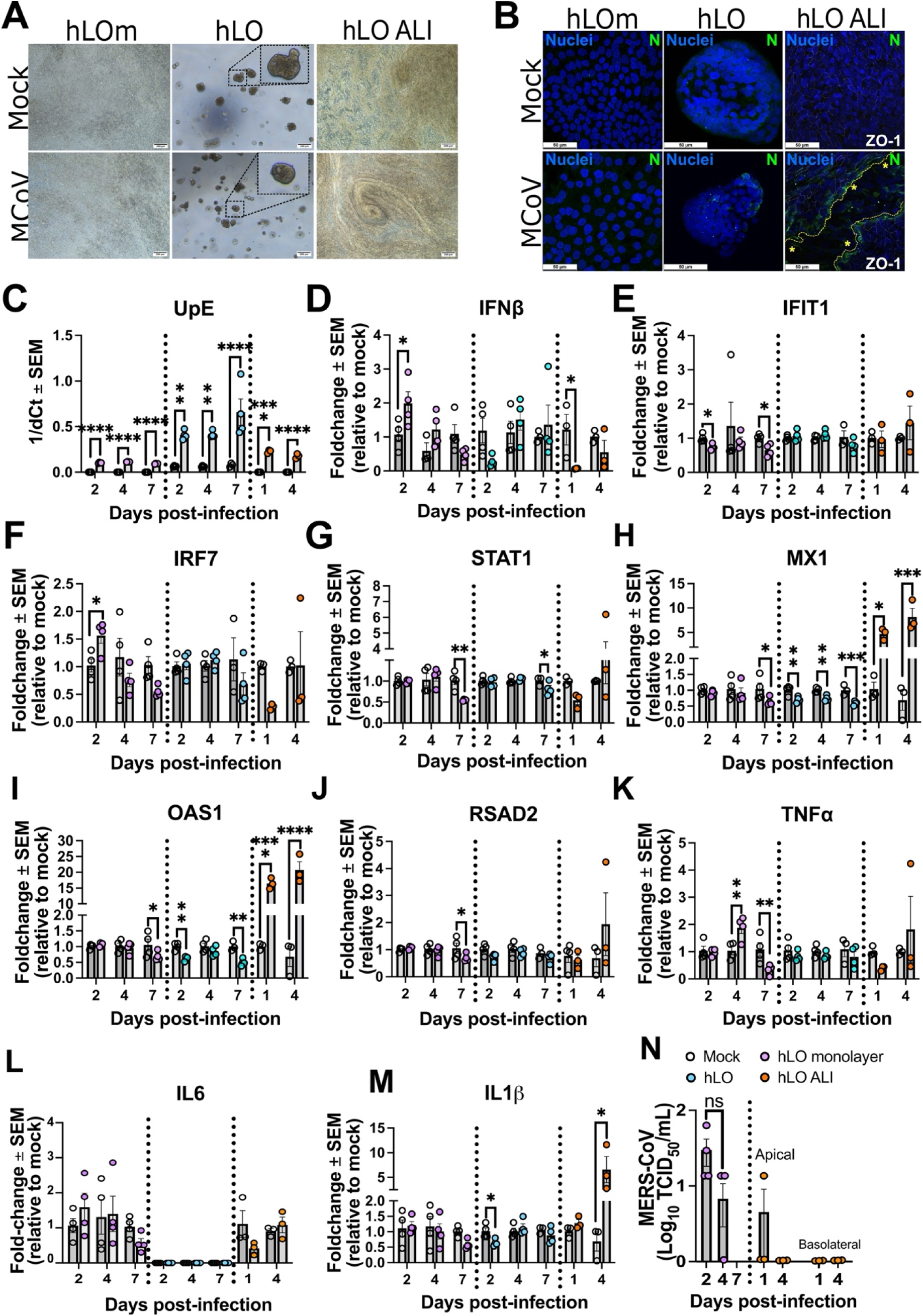
MERS-CoV infectivity in donor-derived lung organoids (hLO), organoid-derived monolayers (hLOm) and ALI (hLO ALI) cultures. (A) Brightfield and (B) confocal imaging of immunostained cells mock infected or infected with MERS-CoV (MCoV) at an MOI of 0.3 at 4 days post infection. Cells were fixed, permeabilized and stained with antibodies against the MERS-CoV nucleoprotein (N) and DAPI (nuclei) and imaged at 63x. Scale bars correspond to 200 µm (A) and 50 µm (B). Dashed lines and asterisks (yellow) delineate areas within the ALI in which ZO-1 was absent, and the culture integrity was disrupted. Total RNA was collected at specified times to determine expression of a viral genomic region upstream the E gene (UpE) (C) and host gene expression of *IFNβ* (D), *IRF7* (E), *IFIT1* (F), *STAT1* (G), *MX1* (H), *OAS1* (I), *RSAD2* (J), *TNF-α* (K), *IL6* (L) and *IL1β* (M) by RT-qPCR. (N) MERS-CoV virus titers in medium supernatant of hLOm, apical and basolateral aspects of hLO ALI, by TCID_50_ assay. The mock infected samples for hLO ALI are the same as in figure 3. For RT-qPCR assays, samples were assayed in technical duplicates, dCt was normalized to GAPDH and virus gene expression was calculated as 1/dCt. Host gene expression was calculated as fold-change relative to mock infected controls using the 2^-ddCt^ method. The mean of three independent samples is presented, error bars are standard error of the mean (SEM). Statistical analysis was performed by One-way ANOVA (C-M) or Student’s T Test (N). *, *P* < 0.05; **, *P* < 0.01; ***, *P* < 0.001; ****, *P* < 0.0001.

Transcripts for antiviral genes like *MX1* (**Fig. 4H**) and *OAS1* (**Fig. 4I**), along with proinflammatory genes like *IL1β* (**Fig. 4M**) were upregulated following infection with MERS-CoV in hLO ALI. Interestingly, these genes were downregulated in hLO and hLOm. Transcripts for *IFNβ* and *IRF7* were upregulated at early time points post infection in hLOm (**Figs. 4D-F**). In hLOs, we noted mostly discrete downregulation of antiviral and proinflammatory gene transcripts. Due to the lower levels of UpE relative to hLO, we performed TCID_50_ assays to determine whether hLOm and hLO ALI cultures released infectious virus progeny into culture media and the apical side of ALIs (**Fig. 4N**). Like SARS-CoV-2, hLOm released infectious virus at Day 2, which decreased over time (**Fig. 4N**). We did not detect infectious virus in apical washes of hLO ALIs or in ALI basolateral medium (**Fig. 4N**). These results show that hLO ALI recapitulated MERS-CoV infection, host response, and pathological tissue damage which was undetectable in hLO and hLOm cultures.

## 3. Discussion

Lung organoids are emerging tools to identify and model respiratory infectious diseases due to their tissue-like cellular heterogeneity and three-dimensional organization (20, 35). Here, we generated donor-derived lung organoids and used them to make two-dimensional cell monolayers (hLOm) and three-dimensional ALI (hLO ALI) cultures. We compared infection kinetics of SARS-CoV-2 and MERS-CoV and associated host responses in these three culture formats. Relative to hLOm and parental hLO, hLO ALI were more responsive to virus infection, allowing for the identification of immune response signatures corresponding to SARS-CoV-2 and MERS-CoV. Our comparative study shows that despite sharing a common origin, each culture format had distinctive virus infection and host response patterns.

An advantage of organoids is that they can recapitulate key features of the respiratory tract. Organoid culture is achieved through growth in gel-like matrix supports to ensure survival and differentiation of stem cells into cell types typical of the tissue of origin. Growth in basement membrane matrix gel prevent exposure to air while promoting growth of organoids as multicellular aggregates typically in the apical-in orientation wherein functional cilia does not develop (36). Despite these disadvantages, organoids from PSC (12, 37), iPSCs, and adult stem cells (11, 38, 39) have been used to generate nasal (40), tracheobronchial (37) and alveolar organoids (41) to model SARS-CoV-2 infection. In contrast, only few studies have used respiratory organoids for MERS-CoV research (42, 43). As reported previously, we found that organoids generated from healthy donor tissue differentiated into cell types typical of tracheobronchial tissue in the apical-in orientation (**Fig. 1**) and were readily infectable by SARS-CoV-2 (**Fig. 3**) and MERS-CoV (**Fig. 4**). However, infecting hLOs involved the recovery of matrix-free organoids followed by virus infection and organoid re-embedding in matrix, which is a complex process that does not facilitate high throughput work. Along with the lack of cilia and exposure to air, these are significant drawbacks for experimental modelling of respiratory diseases using organoids.

Given current challenges, we were prompted to explore the versatility of organoids in two additional culture formats: as cell monolayers (2D) and as ALI cultures (3D). Cell line monolayers composed of homogenous cell populations expressing virus receptors are the cornerstone for studies on SARS-CoV-2 and MERS-CoV. Cell lines like Calu-3 (44–47), Huh-7 (48), MRC-5 (49), and the non-human primate cell line Vero and its derivates (50) have been widely used in coronavirus biology research. We used organoids to produce monolayers and analyzed virus infection kinetics. Unlike cell lines in which viruses cause widespread damage (cytopathic effect) due to quick and efficient replication, monolayers derived from primary organoids were infected poorly by SARS-CoV-2 (Fig. 3) and MERS-CoV (Fig. 4). Transition from organoids into monolayers led to the loss of detectable ACE2 and TMPRSS2 expression levels (Fig. 2H), resulting in a loss of viral infectivity in these cells.

hLOm were found to be immunocompetent against viral dsRNA analogs (Fig. 2A-F) but were unresponsive to bacterial LPS (Fig. 2G), indicating that monolayers do not retain the full breadth of immune response signalling. Human primary tracheobronchial cells upregulate IL6 and IL- 1β when exposed to high doses (10-100µg/mL) of LPS *in vitro* (51, 52), although high LPS dosage has also been linked to loss of viability and subsequent release of inflammatory markers (53, 54). However, features observed in primary cell cultures cannot be extended to hLOm due to differences in culture establishment and conditions such as growth media. In our study, the addition of a single growth factor (Epidermal Growth Factor, EGF) was required to direct monolayer-like growth from dissociated organoid cells. In epithelial cells, LPS engages TLR4 at the cell surface in the canonical pathway, which results in transactivation of the EGF receptor (EGFR) with subsequent NF-κβ activation in LPS-dependent acute lung injury models (55, 56). While additional studies are needed to clarify the mechanisms by which hLOm lose responsiveness to LPS, we surmise that it is possible that EGF occupancy or regulation of EGFR may interfere with LPS-dependent inflammation as it occurs in fibroblasts *in vitro* (57).

Our data suggest that cell monolayers derived from organoids may not be appropriate models for mechanistic studies on pathogen-host interactions and care must be observed about the utility of these models to study antiviral response. In contrast, hLO ALI cultures could be infected with SARS-CoV-2 which led to the production of infectious progeny virions that were released at the apical side of the ALI membrane (Fig. 3). hLO ALI cultures upregulated a diverse array of antiviral transcripts more frequently than parental organoids (Fig. 3D-M). Our study is consistent with previous work highlighting the improved immune response against SARS-CoV-2 in ALI cultures, including those generated from primary bronchial cells (58).

Studies exploring MERS-CoV infection in organoids and ALI cultures are scarce. We investigated both culture formats and found that parental organoids (hLO) and hLO ALI were permissive to MERS-CoV infection (**Fig. 4B-C**). Responses against MERS-CoV were centered on upregulation of antiviral transcripts for *MX1* (**Fig. 4H**) and *OAS1* (**Fig. 4I**) as reported in other *in vitro* studies (59, 60). Interestingly, *MX1* and *OAS1* transcripts were downregulated in parental organoids and hLOm infected with MERS-CoV. Induction of *IFNβ* transcripts was largely absent in hLO and hLO ALI cultures, demonstrating that 3D models can also emulate this aspect of MERS-CoV IFN antagonism (61, 62). MERS-CoV infection induced the upregulation of transcripts for proinflammatory cytokines like *IL-1β* in hLO ALI cultures, but not in the other formats (**Fig. 4L-M)**. Thus, in our studies, hLO ALI models better recapitulated responses that underlie hyperinflammatory syndromes in patients with severe MERS (63) and virus-mediated IFN antagonism.

MERS-CoV induces CPE or cell death in various human and non-human cell lines like Calu3, Huh7, Vero and derivates (50, 62, 64, 64). MERS can induce sloughing of infected cells in ALI cultures made from primary human tracheobronchial cells (65). Instead of sloughing, we identified dissolution of tight junctions at the epithelial barrier of organoid-derived ALIs where plaque-like lesions lined by infected cells had formed (**Fig. 4B**). Disturbances in the epithelial barrier have also been identified in human alveolar tissue infected with MERS-CoV *ex vivo* (66) and in postmortem histopathological exams (67). Our study suggests that organoids at the ALI can recapitulate part of the virus-induced tissue damage and immune responses better than their parental organoids.

Taken together, despite their shared origin, 2D and 3D cell culture formats influenced pathogen kinetics and host immune responses in our studies. These differences between cultures in 2D and 3D cultures must be taken into consideration when selecting models to investigate respiratory diseases. As technology continues to advance, bioengineering approaches like microfluidic systems can help refine current respiratory models for infectious disease research and therapeutic testing. Indeed, our work informs the selection and use of human-derived models compatible with biomimetic approaches for infectious disease research and therapeutic screening. Given the limitations of organoids that typically grow in apical-in orientations, organoids at the ALI are a more reliable and reproducible model for infection and immune response studies that mimic the air-liquid environment of the native lung.

## 4. Materials and methods

### Generation of human lung organoids

This study was approved by the University of Saskatchewan’s human research ethics committee under secondary use of biological material (Permit number: Bio 3570).

Human lung organoids (hLO) were established as described by Sachs (21) from a healthy tissue sample obtained from the upper lobe of a male patient undergoing surgery. Briefly, lung tissue was minced in Advanced DMEM/F-12 (Gibco, cat. 12634010) before single cell dissociation in ACF dissociation solution (StemCell, cat. 05426) for 1.5h at 37°C. Undigested tissue was pelleted, and the cell suspension was strained through a 37µm mesh reversible strainer. Red blood cells were lysed with ACK lysing buffer (Gibco, cat. A1049201) and the remaining cells were embedded in Matrigel (Corning, cat. CACB356231). Matrigel-embedded cells were cultured in complete organoid media supplemented with antibiotic-antimycotic (Gibco, cat. 15240096). hLOs were passaged every 7-10 days by trypsinization and re-embedding into fresh Matrigel. To generate cell monolayers from hLOs (hLOm), dissociated cells were cultured in complete hLO media supplemented with EGF (20 ng/mL). Media was changed every 3-4 days until reaching 80% confluence before passage or used in indicated experiments. Air-liquid interface cultures (hLO-ALI) were generated using single cells from hLOs seeded on transwell inserts (StemCell, cat. 38024) in hLO complete media supplemented with EGF (20 ng/mL) in top and bottom chambers until reaching confluence. Bottom chamber media was replaced with ALI maintenance medium (Pneumacult-ALI, StemCell, cat. 05001) and replaced every two days. The top chamber was exposed to the air interface for 28 days to allow differentiation into mucociliary pseudostratified epithelium. The apical aspect of ALI cultures was washed in 200µL of PBS for 10 minutes at 37°C at least once a week after exposure to air to remove mucus. Basal media was changed every other day until experiments.

### Virus infection

SARS-CoV-2/SB2 clinical isolate (68) and MERS-CoV isolate EMC/2012 were used in these experiments. hLOs were released from Matrigel in ice cold gentle harvesting buffer (Cultrex, cat. 3700-100-01) and infected at an MOI of 20 per hLO (∼0.3 per cell) for 2 hours at 37°C in Advanced DMEM/F-12. hLOs were washed twice to remove unbound virus, re-embedded in Matrigel and cultured in hLO complete media for indicated times. Monolayers were cultured until 80% confluence, washed with PBS and infected at MOI 0.3 for 1 hour at 37°C. Cells were PBS washed and incubated for the indicated times in hLO complete media supplemented with EGF. The apical side of differentiated ALI cultures was infected at MOI 0.3 for 1 hour at 37°C to mimic virus entry through the air interface. Infected ALIs were washed twice and incubated in ALI maintenance medium for indicated times. After indicated times, total RNA was collected using the RNeasy Mini kit (Qiagen, cat 74104) following the manufacturer’s instructions. Work with infectious virus was performed in containment level 3 (CL-3) facilities at VIDO-Intervac, University of Saskatchewan.

### Immunocompetence assays

hLOs, hLOm and hLO-ALI were exposed to LPS 100 ng/mL (Invivogen, cat tlrl-eblps) prepared in complete media for 6h or transfected with poly(I:C) rhodamine (InvivoGen, cat tlrl-picr) at increasing concentrations for 48 hours using lipofectamine 3000 (Invitrogen, cat. L3000015). RNA was extracted with the RNeasy Mini kit (Qiagen, cat 74104) following the manufacturer’s instructions.

### RT-qPCR

cDNA synthesis from ∼500 ng of RNA was generated with the iScript gDNA Clear cDNA Synthesis Kit following the manufacturer’s instructions (BioRad 1725034). cDNA was diluted 1:10 in RNAse free water and used as template for qPCR reactions using Ssoadvanced™ Universal SYBR kit following the manufacturer’s instructions (BioRad 1725274) using selected primers (Table 1). qPCR reactions were performed in a StepOne Real Time PCR System (Applied Biosciences). We normalized Ct values by GAPDH (dCt) and used 1/dCt for viral transcript levels (UpE), and 2^-ddCt^ method for quantitation of host gene expression relative to time-matched mocks or vehicle controls.

**Table 1.**
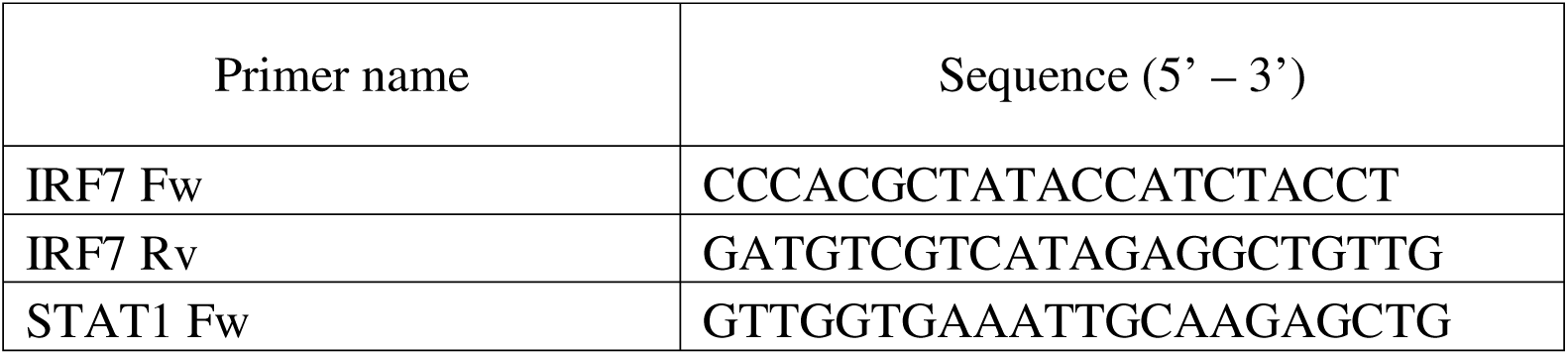
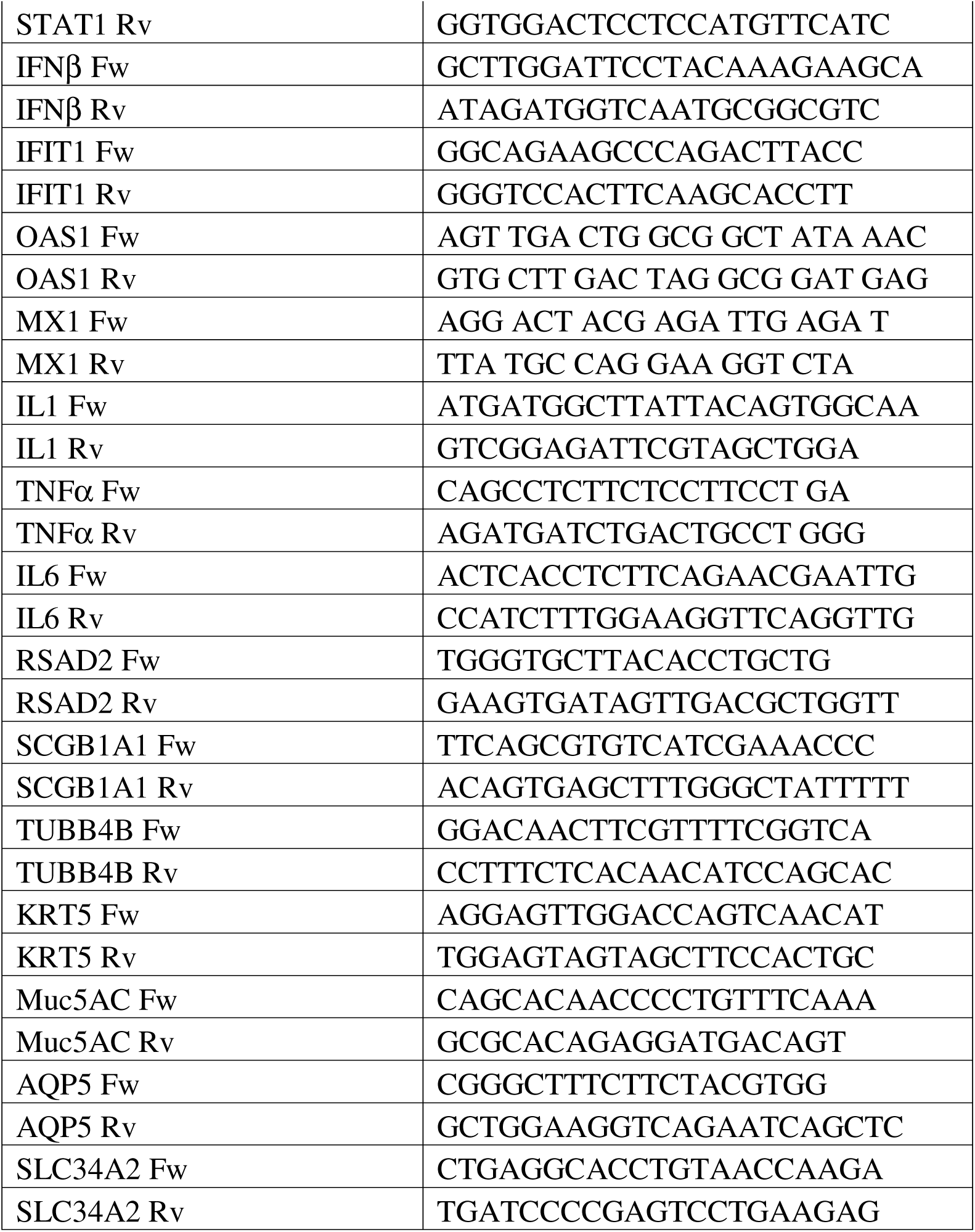
Primers used in this study to determine expression of host genes.

### Immunocytochemistry (ICC)

Matrigel-free hLOs fixed in neutral buffered formalin (NBF) 10% were blocked and permeabilized for 2 hours at room temperature (5% BSA, 1% Triton-X-100, 0.1% Tween-20 in PBS). Incubation with primary antibodies at 1:250 dilution (Table 2) in IF buffer (0.1% BSA, 0.2% Triton-X-100 in PBS) overnight at 4°C. Samples were washed in IF buffer thrice before addition of labeled secondary antibodies (1:3000). Nuclei were counterstained with DAPI (4 μg/mL) prepared in IF buffer, rinsed once in PBS and once in distilled water, and mounted on glass slides using ProLong Gold antifade mountant (Invitrogen, cat. P36930). Monolayers cultured in chambered slides (IBIDI 80806) and ALI cultures were fixed with neutral buffered formalin (NBF) 10% and processed as in ALI. Samples were imaged in Leica TCS SP8.

**Table 2.**
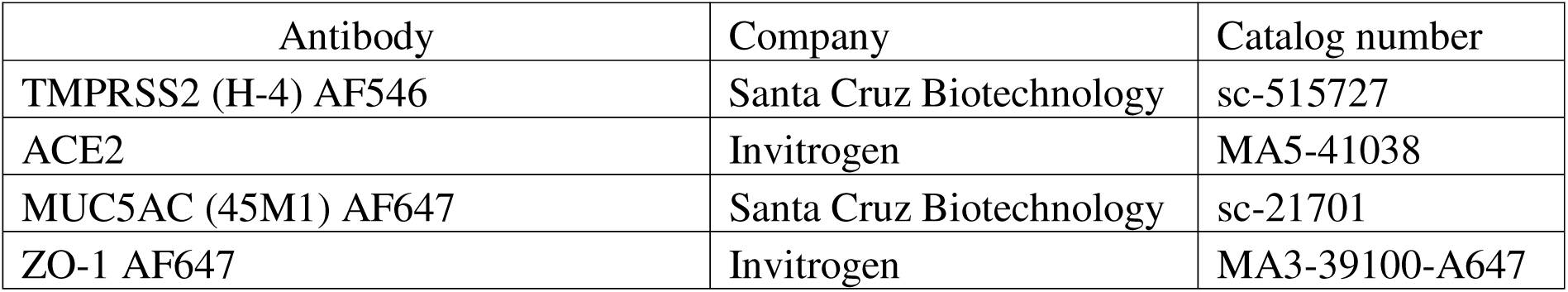
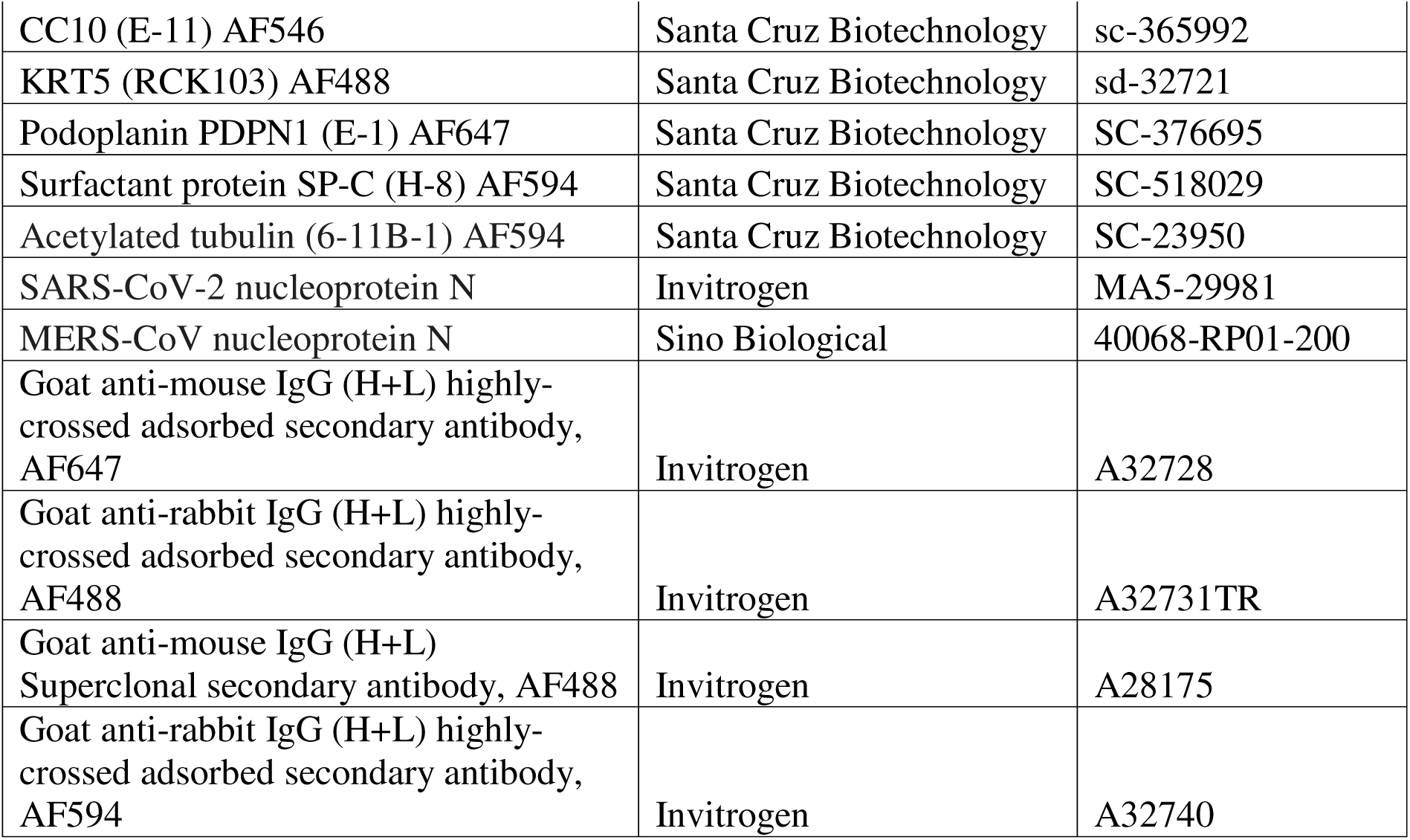
Antibodies used in this study.

### Virus titration (TCID_50_ assays)

Apical washes and basal medium of mock and infected ALI cultures were collected for titration assays. Apical washes were obtained by adding 200uL of PBS to the apical side of ALI cultures and incubating for 10 minutes at room temperature before collection. Supernatant from cell monolayers were also collected. Virus titration was performed as described by us previously. (28, 69, 70). In brief, Vero76 cells were seeded at 2x10^4^ cells per well in 96 well plates and incubated for 24 hours at 37°C in complete media (DMEM supplemented with 10% FBS, 1% penicillin-streptomycin and 1X Glutamax). Vero 76 cells were infected in triplicates with undiluted and tenfold diluted samples prepared in serum free DMEM for 1 hour at 37°C. Inoculum was removed and Vero76 cells were reconstituted with 2% FBS media and incubated at 37°C (5% CO_2_) for up to 5 days. Cells were monitored at 3- and 5-days post infection to determine CPE. Virus titers were calculated according to the Spearman and Karber method as performed previously by us (71).

### Statistics

Statistical analyses were performed in GraphPad Prism (version 10.1.0 (262); www.graphpad.com). RT-qPCR and TCID_50_ data were analyzed by One-way ANOVA or Student’s T Test. Significance values and statistical tests are indicated in the figures and figure legends. *, *P*<0.05; **, *P*< 0.01; ***, *P*<0.001; and ****, *P*<0.0001.

## Supporting information

hLO ALI displayed packed pseudostratified epithelium with presence of beating cilia that actively moved mucus atop the culture

## Acknowledgements

This work was supported by a Saskatchewan Health Research Foundation (SHRF) and Lung SK Solutions - Impact grant (#6185) awarded to A.B. and N.D. K.R.C. was supported by a Living Skies Postdoctoral fellowship awarded by the University of Saskatchewan, Canada. Research within A.B.’s lab was also supported by a Canadian Institutes of Health Research (CIHR) – Institute for Infection and Immunity Early Career Research grant (PTT-192089) and CIHR - Pandemic Preparedness and Health Emergencies Early Career Investigator grant (PEE-183995). Research in ND’s lab was supported by funding from Canadian Institutes of Health Research (ARB-185715 and ARB-192058). VIDO receives operational funding from the Government of Saskatchewan through Innovation Saskatchewan and the Ministry of Agriculture and from the Canada Foundation for Innovation through the Major Science Initiatives Fund. We would like to thank B. Haagmans and R. Fouchier, Erasmus Medical Center, for providing MERS-CoV (isolate hCoV-EMC/2012) and Michelle Gerber for help with organoid culture and maintenance.

